# Amorphous-to-Rodlet Structural Transition Governs the Interfacial Functions of *Aspergillus oryzae* Hydrophobin RolA

**DOI:** 10.64898/2026.02.08.702184

**Authors:** Daiki Ida, Nao Takahashi, Yuki Terauchi, Takumi Tanaka, Akira Yoshimi, Hirotaka Kobayashi, Ken Miyazawa, Masaya Mitsuishi, Hiroshi Yabu, Keietsu Abe

## Abstract

Hydrophobins are low-molecular-weight biosurfactant proteins that coat the cell surface of filamentous fungi, making it hydrophobic and supporting morphogenesis. On conidia, hydrophobins self-assemble to form rod-shaped multimeric structures known as rodlets. Previously, we reported that hydrophobin RolA from the industrial fungus *Aspergillus oryzae* first forms an amorphous film at the air–water interface and then undergoes structural rearrangement to form a densely packed rodlet film. This raised the question of whether the amorphous or the rodlet film is more important for the biological functions of RolA. In this study, to compare the properties of amorphous films with those of rodlet films, we used RolA mutants that had lost the ability to form rodlets and therefore remained in the amorphous state. We found that the rodlet film was more rigid than the amorphous film and had stronger surface activity and a greater capacity to change surface wettability. RolA altered the properties of *A. oryzae* conidia only when it was in the rodlet state. These findings highlight the functional versatility of RolA and show that their dynamic structural transitions directly modulate their function.

## 1 Introduction

Fungal species are thought to be among the most successful taxonomic groups on earth, with more than 3 million species thought to exist (Hawksworth and Lücking R et al., 2017). They are widely distributed and play an important role as decomposers in biogeochemical cycles, because they produce a variety of enzymes that hydrolyze polyesters and polysaccharides (O’Brien et al., 2005, Rytioja et al., 2014; Chen et al., 2013).

Hydrophobins are low-molecular-weight (<20 kDa) amphiphilic proteins conserved in filamentous fungi. Some filamentous fungi use them when they grow in the terrestrial environment and degrade solid polymers (Tanaka et al., 2022). Secreted hydrophobins attach to the cell wall surface and coat it, making it hydrophobic. This surface modification enhances the air-dispersibility of conidia and helps hyphae attach to hydrophobic surfaces such as those of plant leaves (Cai et al., 2020; Talbot et al., 1993). Hydrophobins also contribute to immune evasion by coating the surfaces of pathogenic filamentous fungi, preventing recognition by the host immune system (Aimanianda et al., 2009; Valsecchi et al., 2019). Hydrophobins adsorb hydrophobic solid polymers and then recruit hydrolases, thereby promoting the degradation of these polymers (Takahashi et al., 2005, 2015; Tanaka et al., 2017; Pham et al., 2016). On the basis of such interface-specific functionalities of hydrophobins, various applications have been developed. For example, hydrophobin coating stabilizes nanoparticles in liquid and enables their use in drug delivery systems (Reuter et al., 2017; Valo et al., 2013). Owing to their strong surface activity, hydrophobins have also been used as foaming agents in food and cosmetics (Green et al., 2013). The ability to promote solid polymer degradation has been used for degradation of plastics such as polybutylene succinate-coadipate and polyethylene terephthalate (Takahashi et al., 2005, Puspitasari et al., 2021).

Although hydrophobin amino acid sequences are highly diverse among species, eight Cys residues that form four intramolecular disulfide bridges (Cys1–Cys6, Cys2–Cys5, Cys3–Cys4, Cys7– Cys8) and maintain high conformational stability are well conserved (Kwan et al., 2006). The regions located between Cys3 and Cys4, Cys4 and Cys5, and Cys7 and Cys8 protrude from the molecular surface and are referred to as hydrophobic loops, because they contain many hydrophobic amino acids (Macindoe et al., 2012). Some hydrophobins form rod-shaped polymeric structures called rodlets, which consist of cross β-sheets. Rodlets further assemble densely to form rigid amphiphilic films on the conidial surface (Erwig and Gow et al., 2016; Pham et al., 2016; Terauchi et al., 2020; Valsecchi et al., 2019). Rodlets are highly insoluble and do not depolymerize unless strongly hydrophobic acids such as trifluoroacetic acid or formic acid are applied (Lo et al., 2014; Wessels et al., 1991).

Previously, we analyzed the rodlet formation mechanism of *Aspergillus oryzae* hydrophobin RolA at the air–water interface. We revealed that RolA forms an amorphous monolayer at the interface and then self-assembles into rodlets (Takahashi et al., 2023; Terauchi et al., 2020).

However, it remains unclear which structural state of RolA is responsible for the interfacial functions such as surface wettability modification and high interfacial activity. This uncertainty in the structure–function relationship obscures the molecular mechanism of RolA functionality. In general, interfacial activity of surfactants depends on their interfacial enrichment and molecular assembly structures formed at the interface. Therefore, it is necessary to control the interfacial enrichment and assembly structures of RolA, and subsequently establish an experimental system to evaluate the interfacial properties of each structural state.

In this study, we aimed at elucidating the functional differences between the amorphous and rodlet states of RolA. We hypothesized that the interfacial properties differ between the two states, leading to differences in surface modification ability. To test this hypothesis, we created RolA mutants that are unable to form rodlets and remain in the amorphous state and compared the interfacial structures and properties of the films formed by wild-type (WT) and mutant RolA. RolA-WT formed rodlets at the air–water interface and established robust rodlet film with interfacial characteristics clearly distinct from those of the films formed by mutant RolA. We also investigated the functions of RolA-WT and the mutants on the conidial surface of *A. oryzae* and showed that the self-assembly of RolA controls conidial surface properties. These findings indicate that RolA functions in the rodlet state rather than in the amorphous state.

## 2 Materials and methods

### 2.1 Strains and media

*Aspergillus oryzae* strains used in this study are listed in Table S1. Strains were grown on potato dextrose agar (PDA; BD Difco, Sparks, MD, USA) medium or Czapek–Dox (CD) medium. CD medium contained 1% (w/v) glucose, 0.6% NaNO_3_, 0.052% KCl, 0.152% KH_2_PO_4_, 0.05% MgSO_4_·7H_2_O, 1× trace elements (pH 6.5) (1000× trace elements: 3.6 mM FeSO_4_, 30.7 mM ZnSO_4_, 1.6 mM KH_2_PO_4_, 7 mM MnSO_4_, 0.3 mM Na_2_B_4_O_7_, 0.04 mM (NH_4_)_6_Mo_7_O_24_·4H_2_O), and 2% agar. To complement RolA, malt medium containing 2% malt extract (Thermo Fisher, MA, USA), 1% yeast extract (Thermo Fisher, MA, USA), and 2% agar were used. To transform *A. oryzae*, CD medium containing 0.8 M NaCl was used. To culture the auxotrophic strains, 7 mM sodium hydrogen L(+)-glutamate monohydrate for strain *niaD*^−^, 0.0015% L-methionine for strain *sC*^−^, and 0.01% adenine for strain *adeA*^−^ were added. To construct RolA-complemented and RolA-disrupted plasmids, *Escherichia coli* DH5α Competent Cells (Takara Bio Inc., Shiga, Japan), DynaCompetent Cells JetGiGa DH5α (BioDynamics Laboratory Inc., Tokyo, Japan), and iVEC3 strain (Nozaki and Niki, 2019) were used; cultures were incubated at 37°C in LB broth (Nacalai Tesque, Inc., Kyoto, Japan) containing 100 µg/mL ampicillin.

### 2.2 Protein purification

RolA was purified as described in an earlier report (Terauchi et al. 2020) with modifications. Wild-type RolA (RolA-WT) was purified from the eno-hyp strain, a high-expression strain generated by Takahashi et al. (2015). Single mutants were purified from the eno-hyp L137S and eno-hyp L142S strains, and the double mutant from the eno-hyp L137S/L142S strain generated by Tanaka et al. (2014). These strains express the *rolA* gene under the control of the maltose-inducible enoA142 promoter (P-enoA142). Conidia were inoculated into YPM liquid medium (1% yeast extract, 2% Hipolypepton (SHIOTANI M.S., Hyogo, Japan), and 2% maltose) at a final concentration of 1 × 10^6^ conidia/mL. Cells were cultured at 30°C, 140 rpm, for 48 h (eno-hyp) or 18 h (RolA mutant strains). Each culture was filtered through Miracloth (Merck KGaA, Darmstadt, Germany); the pH of the filtrate was adjusted to 8.5 with 0.1 M Tris (pH 10.5), and electrical conductivity was adjusted to about 1.0 mS/cm with Milli-Q (MQ) water. The adjusted filtrate was applied to a Cellufine Q-500 column (3 × 40 cm; JNC Corp., Tokyo, Japan) equilibrated with 5 mM Tris-HCl buffer (pH 9.0).

RolA was eluted with a 0–0.3 M linear gradient of NaCl. Each fraction was confirmed by sodium dodecyl sulfate–polyacrylamide gel electrophoresis (SDS-PAGE; 3% stacking gel and 17.5% running gel). The fraction containing RolA (13.6 kDa) was dialyzed against 10 mM sodium citrate buffer (pH 4.0) and applied to an SP-Sepharose Fast Flow column (2 × 18 cm; GE Healthcare Japan, Tokyo, Japan) equilibrated with the same sodium citrate buffer. RolA was eluted with a 0.05–0.3 M linear gradient of NaCl. The purified RolA was dialyzed in 10 mM ammonium acetate (pH 7) and lyophilized. All procedures were performed at 4°C. Lyophilized RolA was dissolved in MQ water or 10 mM sodium-acetate buffer (pH 5) before use.

### 2.3 Preparation and modification of SiO_2_ substrates

Hydrophilic and hydrophobic modification of the silicon substrate surface was performed as described in an earlier report (Terauchi et al. 2020). To remove microscopic contaminants from the surface, silicon wafers (p-Si wafers, ≤0.02 Ω cm; Mitsubishi Materials Trading Co., Tokyo, Japan) were ultrasonically cleaned with chloroform, acetone, and 2-propanol for 15 min each. To remove organic contamination of the surface, the wafers were cleaned (each surface for 30 min) with an ultraviolet–ozone cleaner (SKB401Y-01; Sun Energy Co. Ltd., Kanagawa, Japan). At this point, the wafers were hydrophilic owing to the presence of Si-OH groups. To make them hydrophobic, they were incubated overnight in chloroform containing 0.1% *n*-octyltrichlorosilane (Tokyo Chemical Industry Co., Ltd., Tokyo, Japan) at room temperature; they were flushed and dried with N_2_ gas.

### 2.3 Langmuir–Blodgett experiments

RolA Langmuir films were fabricated according to an earlier report (Terauchi et al. 2020) in a USI-TRF3-22 Langmuir–Blodgett trough (USI Co. Ltd., Fukuoka, Japan). A Langmuir–Blodgett trough (100 mm × 360 mm) was filled with 10 mM sodium acetate buffer (pH 5.0), and RolA solution was spread on the buffer surface from a micro-syringe to achieve a RolA amount of 80 µg. To monitor the formation of the Langmuir film, surface pressure (*π*) and average area per molecule (*A*) isotherms were measured at 20.0 ± 0.5°C. *π* was measured by using a Wilhelmy plate attached to a sensitive balance with an accuracy of ± 0.01 mN/m. The compressive modulus (*Cs*^−1^) was defined as:

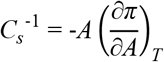

where *T* is the temperature of the aqueous phase. The RolA Langmuir film was compressed at a rate of 3 mm/min and transferred at a constant pressure onto silicon substrates via the vertical lifting method.

### 2.5 Atomic force microscopy

RolA films transferred on silicon substrates were observed under a scanning probe microscope (SPA400; Seiko Instruments Inc., Chiba, Japan) with a cantilever (SI-DF20, Al-coated, f = 136 kHz, C = 16 N/m; Hitachi High-Tech Science Corp., Tokyo, Japan). The sample was scanned in dynamic force mode to obtain the topographic image. The surface irregularities of the sample and the size of the structures formed by RolA were analyzed in observation software (AFM 5000 v. 6.04c, Hitachi High-Tech Science). At least three areas per sample were scanned, and at least three independent experiments were performed per sample.

### 2.6 Measurements of water contact angle

A droplet of MQ water (5 µL) was placed on the transferred RolA films, and the contact angle was measured after 10 s by using a contact angle meter (CA-D; Kyowa Interface Science Co., Ltd, Saitama, Japan). At least three independent experiments were performed for each sample.

### 2.7 Measurement of surface tension

To measure the dynamic surface tension of RolA solution, we used the pendant-drop method (Rotenberg et al., 1983) with a Drop Master 300 surface tension meter (Kyowa Interface Science Co., Ltd., Saitama, Japan) at 20°C. We calculated surface tension values by fitting the droplet profile to the Young–Laplace equation in FAMAS software (Kyowa Interface Science Co., Ltd, Saitama, Japan). Each suspended droplet of RolA solution was prepared inside a screw-capped cuvette to protect it from vibrations caused by air currents and from airborne impurities. To prevent droplet evaporation during the measurement, 10 mM sodium acetate buffer (pH 5.0) was placed in advance at the bottom of the cuvette to produce high humidity around the droplets.

To measure equilibrium surface tension at different RolA concentrations in solution, the measurement time was fixed at 7200 s and the surface tension value at each concentration was measured. It should be noted that surface tension values, especially at low concentrations, may be slightly overestimated. The surface tension values were calculated from protein concentration at the interface by the Gibbs adsorption isotherm as given by the following equation (Wüstneck et al., 1996).

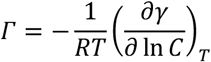

where *R* is the gas constant, *T* is the absolute temperature in K, *γ* is the interfacial tension, and *C* is the RolA concentration. Using this equation, we calculated the *Γ* from the slope of the γ–lnC plot. Minimum are per molecule: *A* was calculated as:

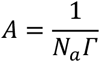

where *N_a_*is the Avogadro constant and *Γ* is surface excess concentration.

### 2.8 Generation of strains with RolA disruption

Sequences of all primers used in the following experiments are listed in Table S2. The *rolA* gene of *A. oryzae* was disrupted by insertion of the adenine auxotrophic marker gene *adeA* from genomic DNA of the *A. oryzae* RIB40 (ATCC-42149) strain (Machida et al., 2005). Polymerase chain reaction (PCR) was used to amplify the *adeA* gene and 1000 bp sequences upstream and downstream of the *rolA* open reading frame (ORF). These three fragments were mixed and used in fusion PCR with the AorolA-top-f and AorolA-bottom-r primer sets. The vector pZErOTM-2 (Thermo Fisher Scientific, Carlsbad, CA, USA) and the insert fragment were ligated using a Mighty Mix DNA Ligation kit (Takara Bio Inc.), and the ligation mixture was transformed into *E. coli* iVEC3 strain (Nozaki and Niki, 2019). The plasmid pZErO-2-ΔrolA was transformed into the *A. oryzae adeA*^−^ strain 15 by the protoplast–PEG method (Gomi et al., 1987; Mizutani et al., 2004). Disruption of the *rolA* gene in the transformants was confirmed by genomic PCR with primer sets (1) AoRolA-delta-L-chk-f2, AnAdeA-chk-r2, (2) AnAdeA-chk-f2, AoRolA-delta-R-chk-r2, and (3) AoRolA-chk-f, AoRolA-chk-r. Genomic PCR was performed using SapphireAmp Fast PCR Master Mix (Takara Bio Inc.) under the following conditions: 1 cycle of 94°C for 1 min, 30 cycles of 98°C for 5 s, 58°C for 5 s and 72°C for 30 s, 1 cycle of 72°C for 5 min. Uninucleate conidia were then purified by passing the conidial suspension at least twice through an Isopore membrane filter (Millipore, Burlington, MA, USA) with a pore size of 5 µm.

### 2.9 Generation of RolA-complemented strains

Plasmids used to generate RolA complementation strains were produced by In-Fusion cloning (Figure S1). The plasmid pNEN142-AorolA WT contains the RolA-WT ORF (557 bp) (Takahashi et al., 2015) but not the P-enoA142 promoter (Tsuboi et al., 2005). Inverse PCR with the In-Fusion Inverse PCR primer set was used to amplify this plasmid. PCR with the In-Fusion genome Fw, Rv primer set was used to amplify a region 1000 bp upstream of the *rolA* ORF from the genomic DNA of the RIB40 strain. Both fragments were linked with an In-Fusion HD Cloning Kit (Takara Bio Inc.) and transformed into *E. coli* DynaCompetent Cells JetGiGa DH5α (BioDynamics Laboratory Inc.). A PrimeSTAR Mutagenesis Basal Kit (Takara Bio Inc.) and the L137S Fw, Rv and L142S Fw, R primer sets were used to introduce the single mutations L137S and L142S, respectively, into the RolA ORF of RIB40p-AorolA WT by site-directed mutagenesis. To create the RIB40p-AorolA L137S/L142S plasmid with a double mutation, the L137S mutation was introduced into RIB40p-AorolA L142S with the L137S/L142S Fw, Rv primer set. The plasmids were linearized with restriction enzyme *Mun*Ι (Takara Bio Inc.) and transformed into the *A. oryzae* Δ*rolA* strain by the protoplast–PEG method. To select candidate RolA-complemented strains, genomic PCR with the primers In-Fusion genome Fw (binding to the RolA promoter from the RIB40 strain) and pNEN142-enoA-RolA Rv (binding within the *niaD* region) was performed using EmeraldAmp PCR Master Mix (Takara Bio Inc.) using 1 cycle of 94°C for 2 min; 30 cycles of 98°C for 10 s, 55°C for 30 s and 72°C for 2 min; and 1 cycle of 72°C for 5 min. The PCR products were separated in 0.7% agarose gel and stained with ethidium bromide. Single bands of interest were cut out, and DNA was extracted using a NucleoSpin Gel and PCR Clean-up kit (Takara Bio Inc.). To confirm the presence of mutations in the *rolA* ORF in genomic DNA, we used the DNA sequencing service of Eurofins Genomics (Ebersberg, Germany). Uninucleate conidia were purified as in 2.8.

### 2.10 Scanning electron microscopy

A PDA agar block was sandwiched between sterile glass plate, inoculated with the conidia of *A. oryzae* on the lateral surface of the agar block and incubated for 5 days. Then, the upper glass was removed, the bottom glass with the specimen was placed in a 15mmφ*8mmH mesh basket. The specimen in the basket was fixed in a beaker with 2.5% glutaraldehyde and 2% paraformaldehyde in 0.1 M phosphate buffer (pH 7.4), dehydrated through 50% acetone for 1 h followed by an acetone series (70%, 80%, 90%, 95%, 100%) for 30 min, critical point dried, and coated with osmium. The conidial surface was observed under a Hitachi Regulus 8220 field emission scanning electron microscope (Hitachi, Tokyo, Japan) at an acceleration voltage of 1.0 kV.

### 2.11 Determination of conidial surface hydrophobicity

Hydrophobicity of the conidial surface was determined by microbial adhesion to hydrocarbons (MATH) assay (Holder et al., 2007; Zhang et al., 2011). RolA-complemented strains were grown on PDA for 7 days. Conidia were harvested and suspended in PUM buffer (2.22% K_2_HPO_4_·4H_2_O, 0.726% KH_2_PO_4_·4H_2_O, 0.18% urea, 0.02% MgSO_4_·7H_2_O, pH 7.1). Each suspension was adjusted to absorbance at 470 nm (*A*_470_) ≈ 0.4, and 3-mL aliquots were dispensed into 5-mL volumetric glass vials. Hexadecane (300 µL) was added to each vial. The vials were vortexed three times for 30 s and allowed to stand at room temperature for 15 min; the hexadecane phase was then carefully removed and discarded. Vials were cooled to 4°C, any residual solidified hexadecane was removed, and the vials were returned to room temperature. *A*_470_ in the resultant suspension were measured. The hydrophobic index (H.I.) was calculated from *A*_470_ as:

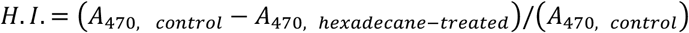

## 3 Results

### 3.1 Rodlet formation at the air–water interface

Sequence alignment of RolA with RodA from *Aspergillus fumigatus* and EAS from *Neurospora crassa* revealed that the residues involved in rodlet formation in RodA (L145) (Valsecchi et al., 2019) and EAS (F98) (Macindoe et al., 2012) corresponded to L137 in RolA, located within the Cys7–Cys8 loop (Figure S2). Our previous study suggested that the self-assembly of RolA is driven by hydrophobic interactions (Takahashi et al., 2023); therefore, it seemed highly likely that this hydrophobic loop contributes to rodlet formation. To reduce its hydrophobicity, we replaced L137, L142, or both with serines and expected that rodlet formation would be reduced in the resulting mutants L137S, L142S, and L137S/L142S.

To evaluate rodlet formation by these mutants, we used the Langmuir–Blodgett method. Consistent with our previous study (Terauchi et al., 2020), the *π* of RolA-WT first gradually increased with decreasing *A* and then transiently dropped to approximately 15 mN/m (Figure 1A), indicating that the protein underwent self-assembly; this structural transition was also evidenced by a *Cs*^−1^ minimum at similar *A* values (Figure 1B). With a further decrease in *A*, the *π* values increased steeply and reached a collapse pressure of about 70 mN/m. In L142S, *π* gradually increased, almost reached a plateau at about 19 mN/m without the transient decrease observed in the WT, and eventually also increased to about 70 mN/m. RolA L137S and L137S/L142S showed a slower onset of the increase in *π* with decreasing *A* and the film collapsed at around 30 mN/m.

**Figure 1.**
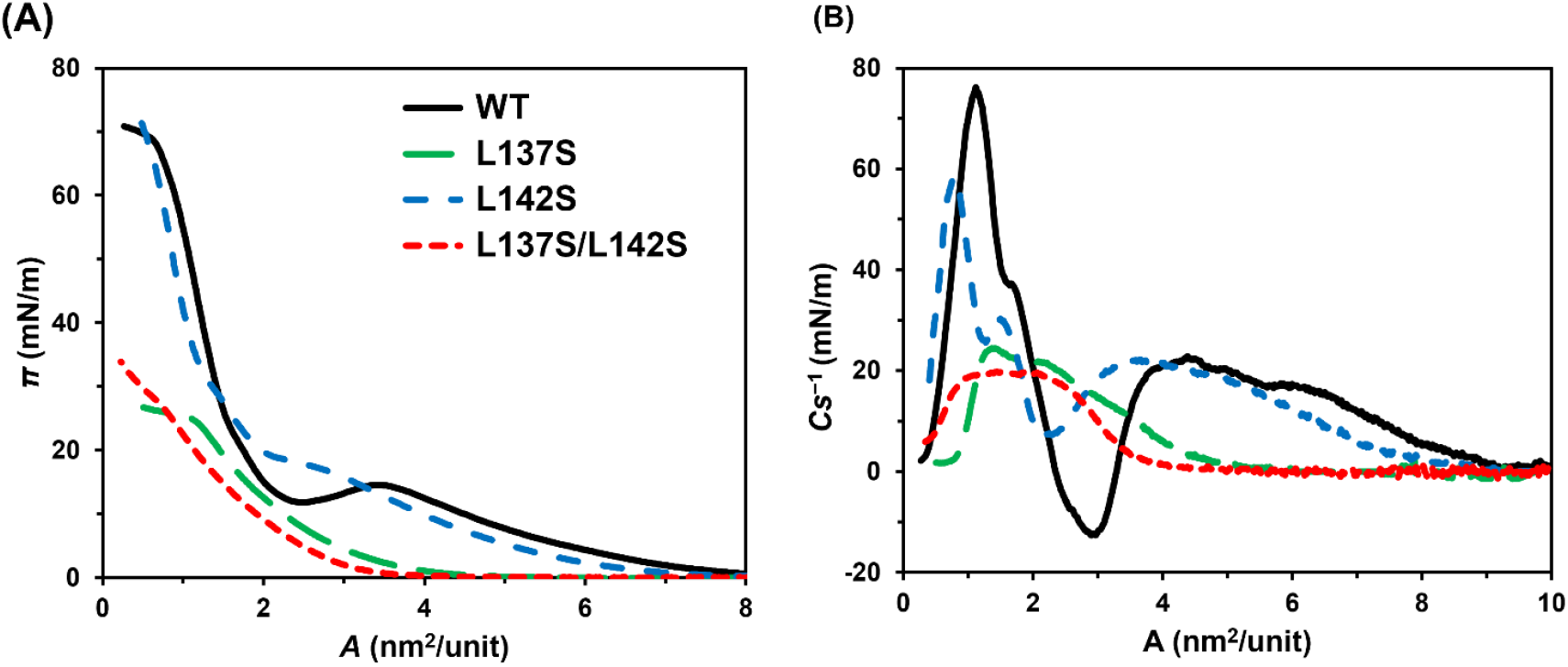
Isotherm profiles of Langmuir films of wild-type (WT) and mutant RolA(L137S, -L142S, - L137S/L142S).

To analyze the structures at the air–water interface, RolA films were transferred onto the hydrophilic silicon substrate, and the RolA structures exposed to the air were observed by AFM (Figure 2), and rodlet height, short axis, and long axis were measured (Table S3). In the WT, consistent with the AFM images in a previous report (Terauchi et al. 2020), a continuous monolayer was observed on the substrate at 10 mN/m and dense rodlet films were observed at 20 and 30 mN/m (Figure 2A). In L137S, a continuous monolayer at 10 mN/m was similar to that in the WT, but markedly fewer rodlets were observed at 20 mN/m than in the WT (Figure 2B). In L142S, spherical structures were found within uniform monolayers at 10 and 20 mN/m, and dense rodlet films, as in the WT, were observed at 30 mN/m (Figure 2C). The size of rodlets formed by the two single mutants was similar to that of the WT rodlets (Table S3). In L137S/L142S, continuous monolayers were observed at 10 and 20 mN/m similar to those in the WT at 10 mN/m, but this mutant did not form rodlets at all (Figure 2D).

**Figure 2.**
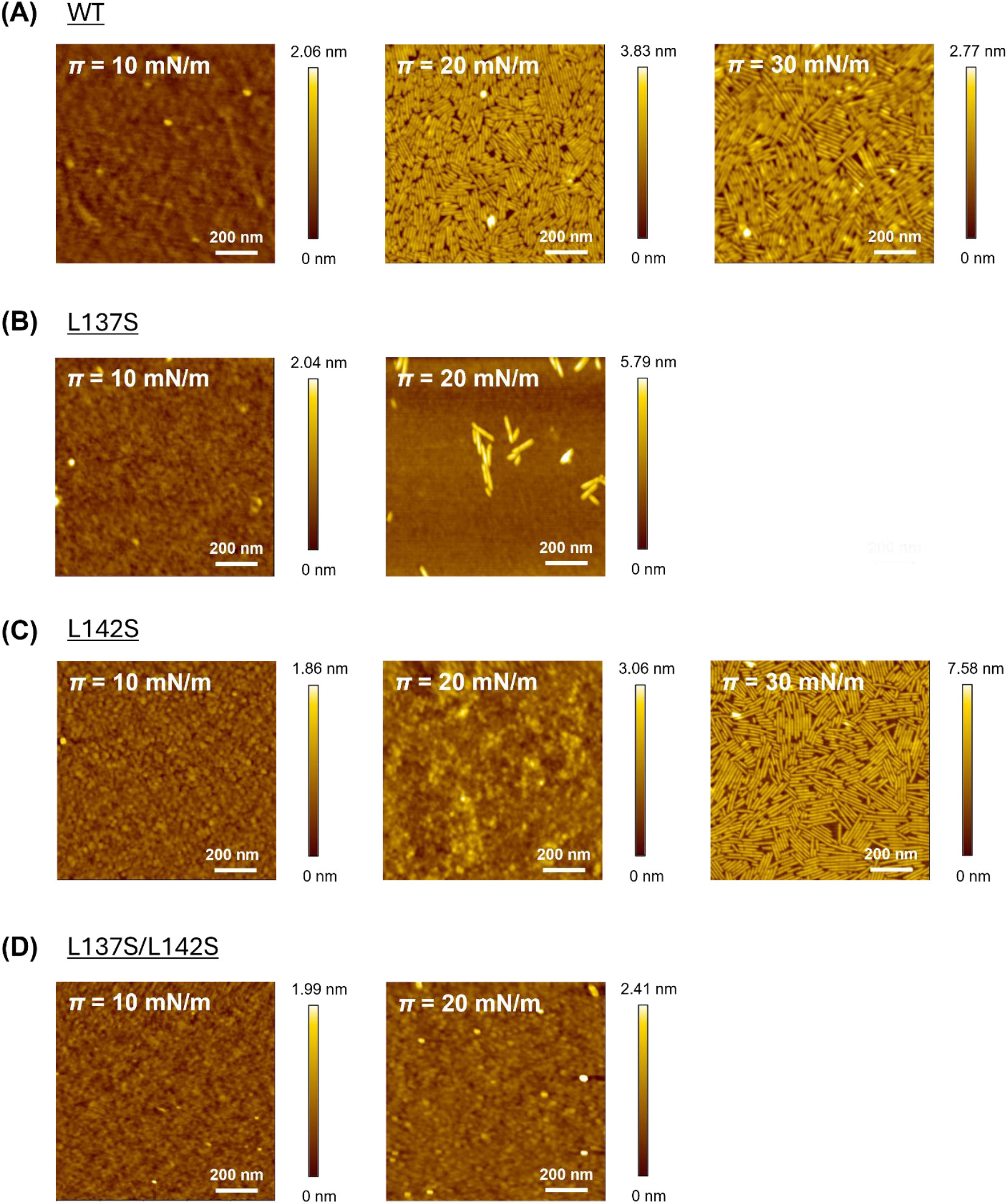
AFM topography images and height profiles of Langmuir films of WT RolA and its mutants. Films were transferred on hydrophilic silicon substrate at the indicated values of *π*. Image size, 1 μm × 1 μm.

The RolA films were also transferred onto the hydrophobically modified substrate and the RolA structures exposed to the liquid phase were observed by AFM. Rodlet formation was observed in the WT but was suppressed in the mutants (Figure S3, Table S3). These data confirm that the Cys7–Cys8 loop of RolA is crucial for rodlet formation, as the RolA mutants remained in the amorphous film state without progression of self-assembly.

### 3.2 Wettability modulation

To assess the ability of RolA and its mutants to change substrate wettability, we transferred RolA films onto the hydrophilic silicon substrate, dropped MQ water onto the films and bare substrate, and compared water contact angles (Table 1). The water contact angle value of amorphous RolA-WT film was higher by about 30° than that of the bare substrate, whereas that of the rodlet film of RolA-WT was higher by about 60°–70°. The contact angle of amorphous film of RolA-L142S was higher by about 30° than that of the bare substrate, whereas that of its rodlet film was higher by about 65°. Since RolA-L137S and RolA-L137S/L142S formed fewer or no rodlets at any surface pressure, the contact angle was higher than that of the bare substrate by only about 20°–30°.

**Table 1.**
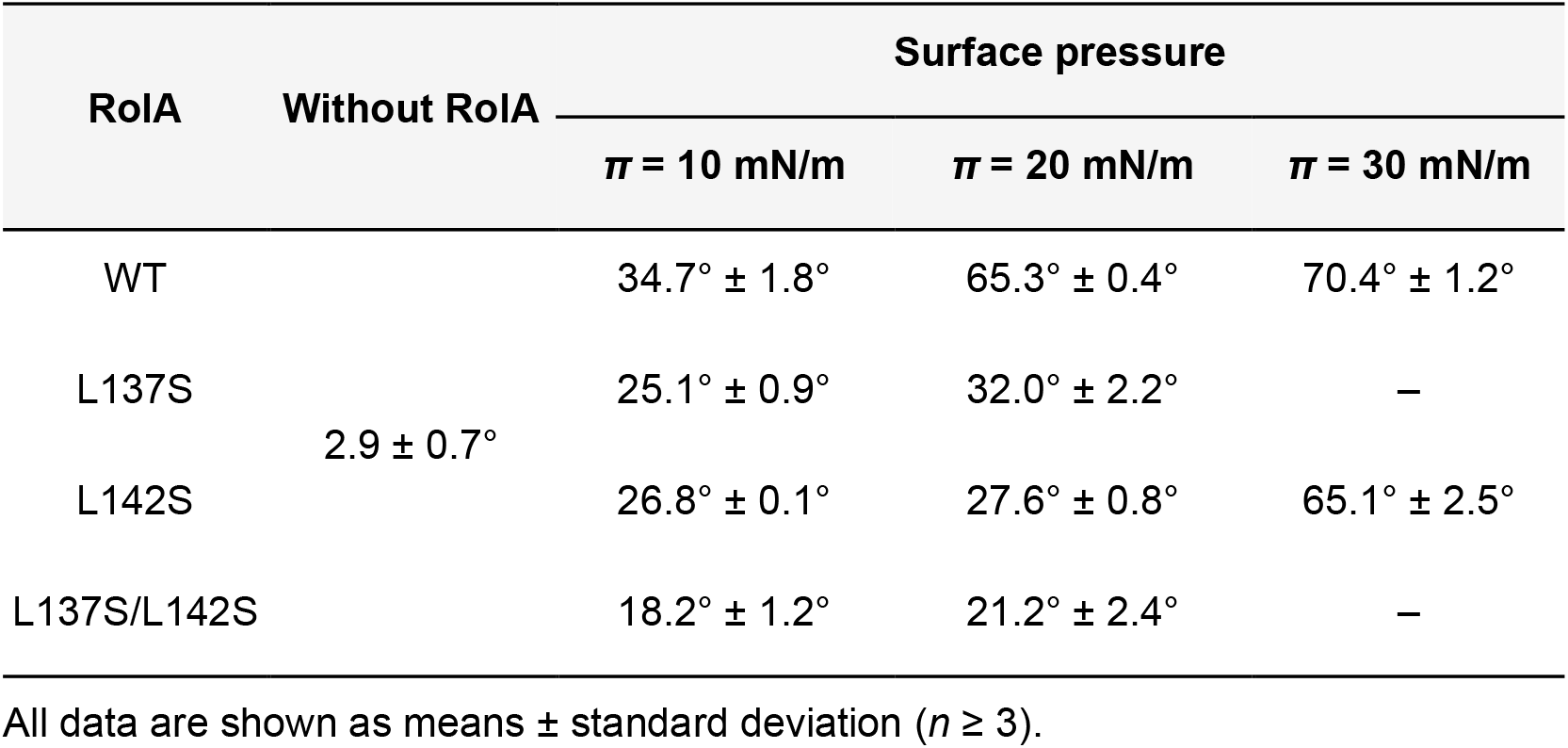
Water contact angles of RolA Langmuir films transferred on the hydrophilic substrate.

We also measured contact angles of RolA films transferred onto the hydrophobic substrate (Table S4). Similarly, rodlet films showed strong surface modification. These results indicate that RolA in the amorphous state does not considerably alter substrate wettability, whereas RolA in the rodlet state does.

### 3.3 Surface activity of amorphous and rodlet films

To examine the effects of the transition from the amorphous state to the rodlet state on the surface activity of RolA, we measured dynamic surface tension using the pendant drop method (Figure 3A). In RolA-WT, surface tension rapidly decreased to 53 mN/m immediately after the start of the measurement, transiently stabilized and then further decreased to 30 mN/m, eventually reaching a plateau. In all the mutants, the surface tension also decreased rapidly after the start of the measurements to about 53 mN/m, and then decreased slowly, reaching about 45 mN/m.

**Figure 3.**
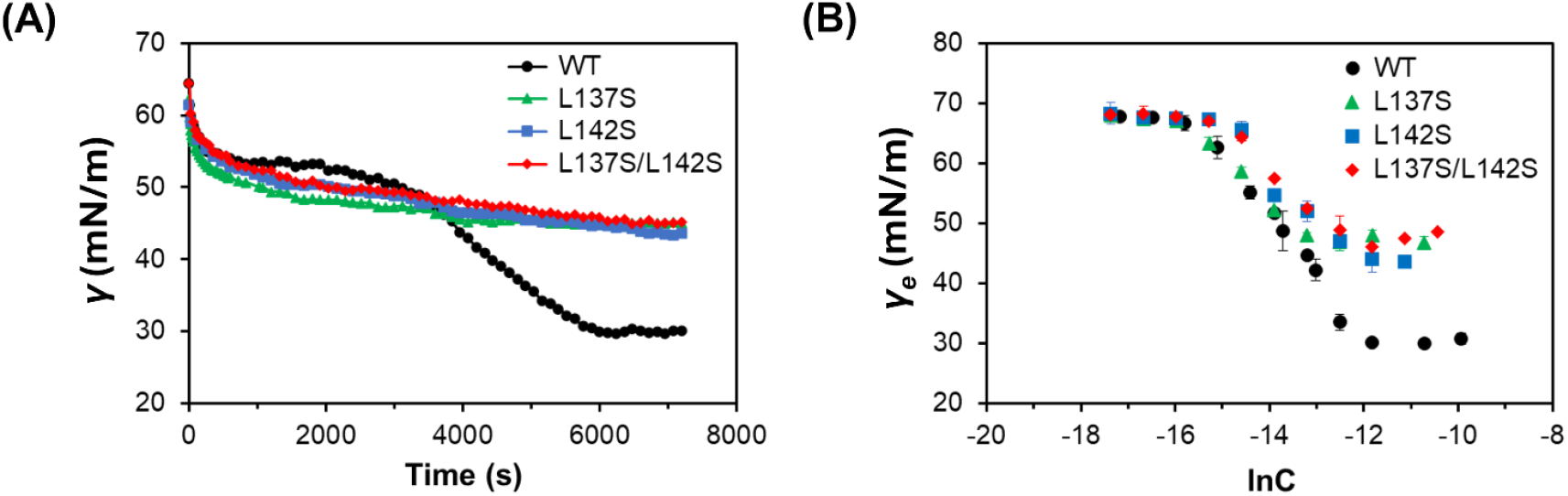
Surface activity of RolA-WT and its mutants. (A) Dynamic surface tension vs. time for RolA at 100 μg/ml. (B) Equilibrium surface tension (*γ*) vs. concentration (ln*C*). Error bars, standard deviation (n = 3).

To investigate the effects of mutations on the amount of adsorbed RolA at the air–water interface, we measured equilibrium surface tension at different RolA concentrations in solution (Figure 3B).

The surface excess concentration and minimum area per molecule are shown in Table 2. The calculated values of *Γ_max_* for the mutants were smaller than that of RolA-WT, and those of *A_min_* were higher than that of RolA-WT. Although the adsorption of proteins may differ from ideal Langmuir adsorption (Rabe et al., 2011), these results demonstrated that RolA-WT undergoes two step changes in interfacial tension, which lead to a markedly higher adsorption and a more densely packed state than in the mutants.

**Table 2.** Parameters derived from the *γ*–ln*C* plot.

| **RoIA** | **$\Gamma_{max} (\times 10^{-11})$ (mol/cm<sup>2</sup>)<sup>a</sup>** | **$A_{min}$ (nm<sup>2</sup>/unit)<sup>b</sup>** |
| --- | --- | --- |
| **WT** | 40.0 | 0.42 |
| **L137S** | 25.4 | 0.65 |
| **L142S** | 34.1 | 0.49 |
| **L137S/L142S** | 26.0 | 0.64 |

<sup>a</sup> $\Gamma_{max}$ : maximum surface excess concentration.
<sup>b</sup> $A_{min}$ : minimum area per molecule.

### 3.4 Qualitative assessment of RolA film stiffness

We checked the stiffness of RolA films formed at the air–water interface (Figure 4). After 7200 s, the droplets of RolA-WT had a pear shape and were constricted near the nozzle (Figure 4A), whereas those of RolA mutants did not undergo such clear deformation (Figure 4B–D). When droplet volume was gradually reduced after dynamic surface tension measurement, the RolA-WT and RolA-L142S droplets became distorted and subsequently formed numerous fine wrinkles around the nozzle (Figure 4A, C). This phenomenon, known as buckling, occurs when an elastic film is formed at the interface (Yamasaki et al., 2016A, 2016B). We defined droplet volume at a certain time point as V and the initial volume as V_0_; buckling was observed at V/V_0_ = 0.66 in RolA-WT and at V/V_0_ = 0.14 in RolA-L142S (Figure S4). No buckling was detected in RolA-L137S or L137S/L142S, even when the droplets shrank (Figure 4B, 4D). Droplets of RolA solutions on a hydrophobic substrate were incubated under humid conditions at room temperature. Consistent with a report that RolA-WT actively self-assembles to form rodlets at the droplet surface (Takahashi et al., 2023), it formed a rigid rodlet film at the droplet surface, resulting in flat drops at 24 h (Figure S5). In RolA-L137S and -L142S, flat drops were formed at 72 h and 48 h, respectively, consistent with suppressed self-assembly. In RolA-L137S/L142S, no flat drops were observed until the drop had fully evaporated, consistent with the lack of self-assembly.

**Figure 4.**
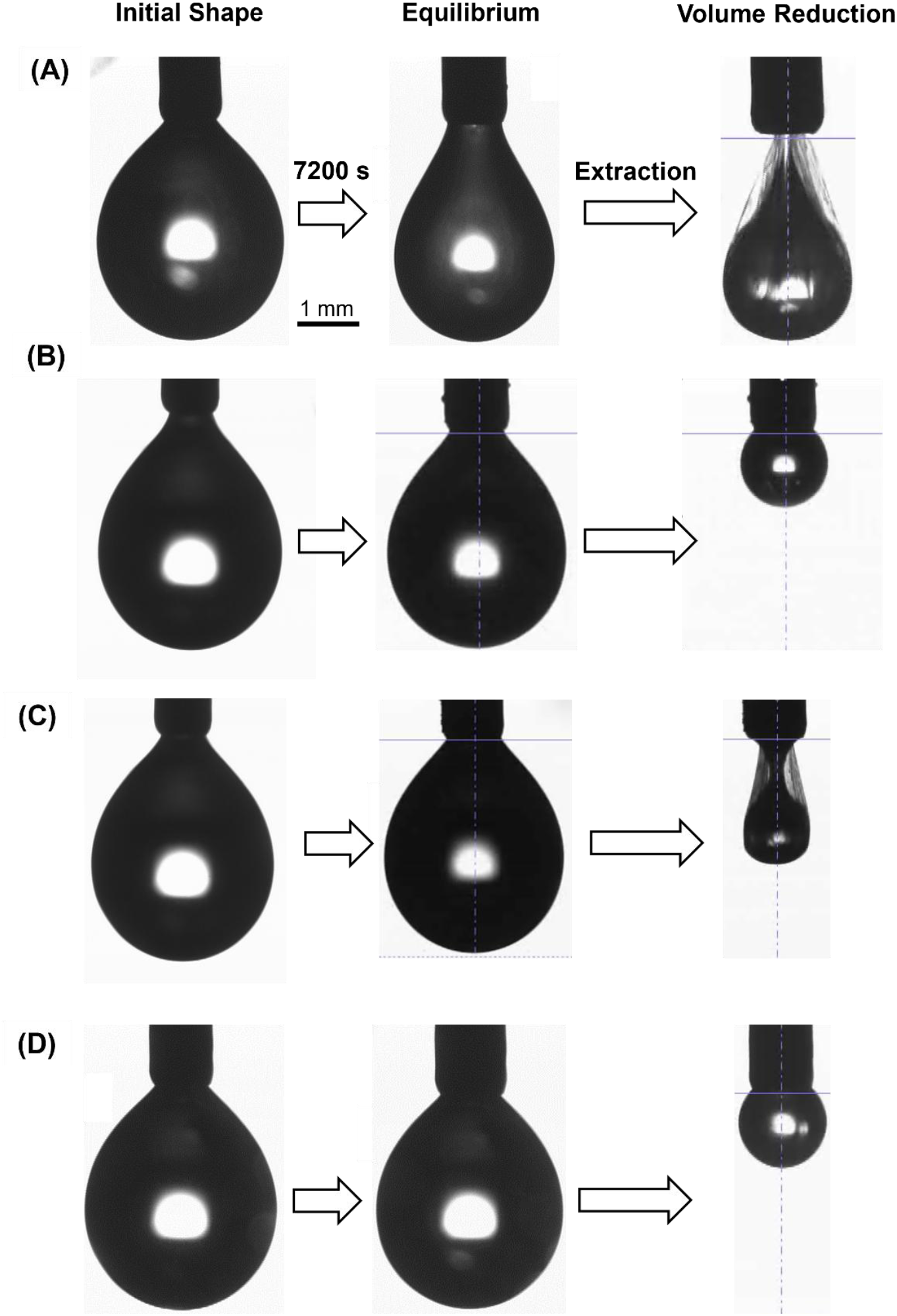
Droplet deformation by RolA films formed at droplet surfaces. Droplet images were taken before and after dynamic surface tension measurements and subsequent extraction of RolA solution for RolA (A) WT, (B) L137S, (C) L142S, and (D) L137S/L142S.

### 3.5 Phenotypic analysis of *A. oryzae* conidia

We constructed RolA-complemented strains in which the expression of *rolA* ORFs encoding WT or mutants was driven by the native *rolA* promoter from *A. oryzae* RIB40 and used SEM to analyze the effects of RolA self-assembly on the surface properties of *A. oryzae* conidia (Figure S5). In the control *adeA*^+^ and RolA-WT^+^ strains, high-density rodlet layers were observed on the conidial surface (Figure 5A, C), whereas no rodlet layer was present in the Δ*rolA* strain (Figure 5B) or in either the single or double mutants (Figure 5D–F). On PDA plates, blackened colonies were observed in the Δ*rolA* strain and the rodlet-defective mutant strains but not in the *adeA*^+^ or RolA-WT^+^ strains (Figure S6). In MATH assay, the H.I. values were approximately 0.7 for the *adeA*^+^ and RolA-WT^+^ strains, and 0.4 to 0.5 for single and ccs (Figure 5G). The differences between the *adeA*^+^ and RolA-WT^+^ and double mutant were statistically significant, indicating that conidial hydrophobicity is reduced in the mutant strains.

**Figure 5.**
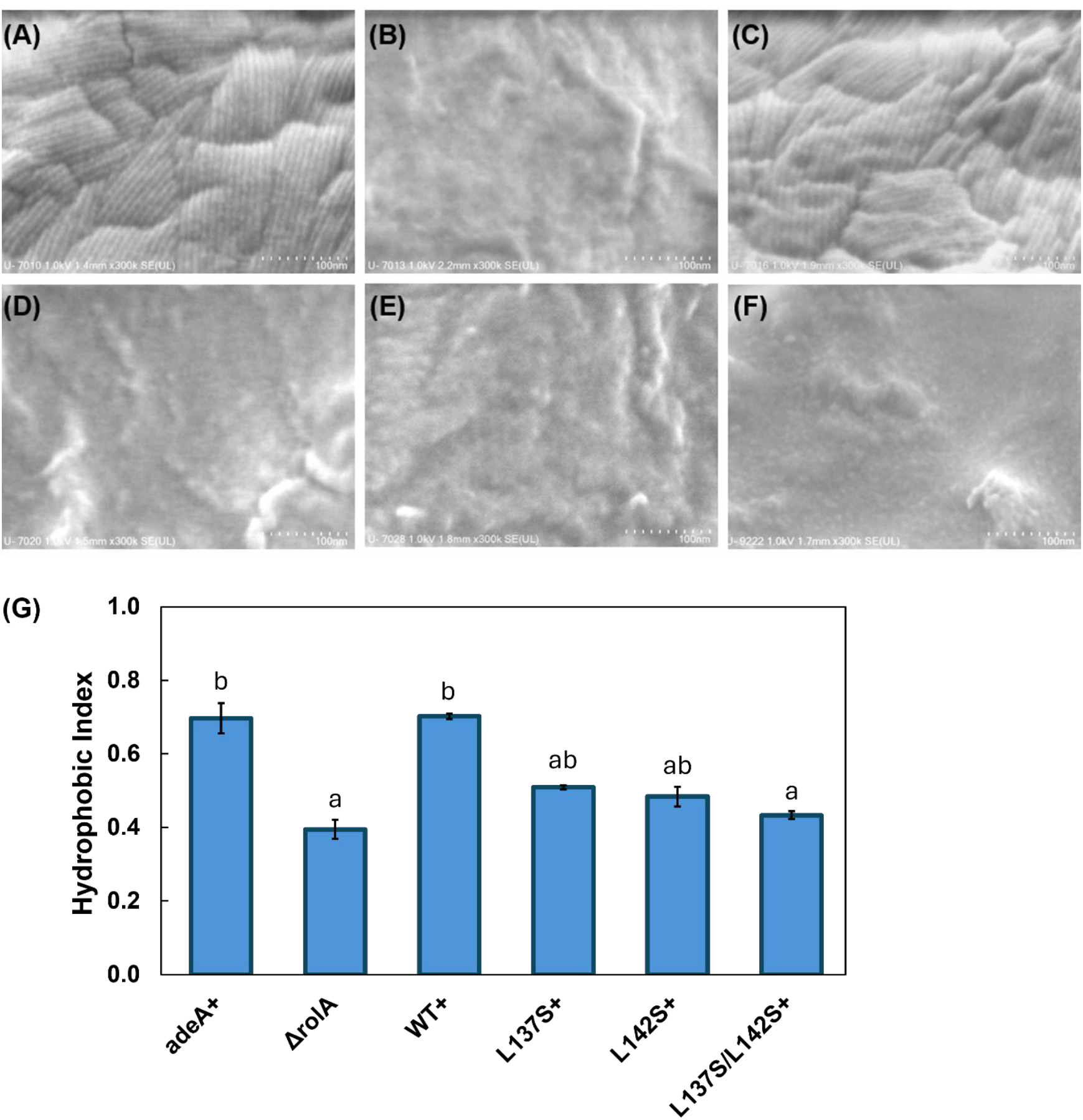
Phenotypic characterization of *A. oryzae* RolA-complemented strains. (A–F) Scanning electron micrographs of the conidial surface of the (A) *adeA_+_*, (B) *ΔrolA*, (C) RolA-WT+, (D) RolA-L137S+, (E) RolA-L142S+, and (F) RolA-L137S/L142S+ strains. (G) Cell surface hydrophobicity index of the conidia of each strain determined by MATH assay. Different letters denote significant differences in Tukey’s test (*P* < 0.05). Error bars, standard deviations (*n* = 3).

## 4 Discussion

In this study, we elucidated the differences between the two distinct structural states of RolA formed at interfaces: the amorphous and rodlet states. Amino acid residues within the Cys7–Cys8 loop were predicted to be involved in rodlet formation (Figure S2). We substituted L137 and L142 within this loop with serines (RolA-L137S, -L142S, -L137S/L142S). In all these mutants, rodlet formation was suppressed to a different extent (Figures 1, 2, S3). Rodlet formation ability decreased in the following order: RolA-WT > L142S > L137S > L137S/L142S. Our previous study suggested that the self-assembly of RolA is driven by hydrophobic interactions (Takahashi et al., 2023). The present findings that hydrophobic Leu residues are crucial for self-assembly support the importance of hydrophobic interactions in RolA self-assembly.

When RolA films created by the Langmuir–Blodgett method were transferred onto hydrophilic and hydrophobic substrates, rodlet films (RolA-WT at 20 and 30 mN/m, RolA-L142S at 30 mN/m) drastically altered substrate wettability, whereas the effect of amorphous films (RolA-WT at 10 mN/m, RolA-L137S at 10 and 20 mN/m, L142S at 10 and 20 mN/m, L137S/L142S at 10 and 20 mN/m) was much weaker (Table 1). These results clearly indicate that surface wettability is modified by RolA to a greater extent when rodlets are formed. However, it should be noted that the amount of RolA Langmuir film transfer onto the substrate requires careful consideration. Defining the transfer ratio as the decrease in monolayer area during transfer divided by the dipped substrate area, the estimated ratios for hydrophilic substrates (Figure 2) are 0.91 (RolA-WT, 30 mN/m), 1.0 (RolA-L142S, 30 mN/m), 1.44 (RolA-L137S, 20 mN/m), and 1.1 (RolA-L137S/L142S, 20 mN/m), whereas those for hydrophobic substrates are 0.7 (RolA-WT, 30 mN/m), 0.5 (RolA-L142S, 30 mN/m), 0.74 (RolA-L137S, 20 mN/m), and 0.83 (RolA-L137S/L142S, 20 mN/m). Because, particularly, hydrophobic substrates are immersed into the trough from air (Terauchi et al., 2020), the RolA film-coated substrate passes through the buffer and the air-buffer interface again during lift-up process, which may cause the film to detach in part. In HGFI from *Grifola frondosa*, the rodlet-forming WT and a rodlet-deficient mutant modify the wettability of mica and glass surfaces to a similar extent (Wang et al., 2017). These findings suggest that the mechanisms of surface modification may differ among hydrophobins, reflecting the diversity in how hydrophobins exert their functions.

Interfacial tension differed between RolA amorphous and rodlet films. RolA-WT had a two-step decrease in interfacial tension, whereas RolA mutants had only a single-step decrease (Figure 3A). This suggests that RolA first forms an amorphous film at the air–water interface, reaches the first equilibrium, and subsequently undergoes self-assembly into rodlets, which induces a further decrease in interfacial tension. Such two-step interfacial tension reduction is not observed in typical low-molecular-weight surfactants (e.g., sodium dodecyl sulfate and Triton X-100, etc.) and can be regarded as a unique feature arising from the combination of the strong surface activity and self-assembly capability of RolA. Estimation of the surface excess concentration and molecular area occupied by RolA-WT and the mutants (Table 2) indicated that RolA-WT formed a more densely packed film structure at the air–water interface when the rodlet state was compared with the mutants of amorphous state. These findings suggest that rodlet formation is essential for RolA to form a dense film structure, which cannot be achieved in the amorphous state, and to have high surface activity.

To better understand the properties of RolA films at the air–water interface, we also analyzed their stiffness after interfacial tension measurements. When a pendant drop of RolA-WT was compressed, buckling was observed (Figure 4A). In contrast, no buckling was detected in RolA-L137S and -L137S/L142S (Figure 4B, D), which do not form rodlets, suggesting that buckling occurs only when a rigid rodlet film is formed on the droplet surface. Buckling was observed later in RolA-L142S than in RolA-WT (Figures 4C and S3). Since RolA-L142S did form rodlet film at high surface pressure in Langmuir–Blodgett experiments (Figure 2C), this reduced rodlet formation ability of this variant likely accounts for the observed buckling. The absence of a two-step decrease in interfacial tension in RolA-L142S during measurements of dynamic surface tension was probably due to insufficient rodlet formation so that the amount of rodlet at the air–water interface did not reach the detection threshold. Differences in flat drop formation among RolA-WT and mutants (Figure S5) were also consistent with their rodlet formation ability, further supporting the notion that droplet deformation occurs when rigid rodlet film is formed at the air–water interface. The formation of highly elastic films at the air–water interface by assembly in other proteins has also been reported. For example, pendant drop buckling has been observed in hydrophobin HFBI of *Trichoderma reesei* (Yamasaki et al., 2016A), chaplin secreted by *Streptomyces coelicolor* (Claessen et al., 2003), and amphiphilic protein BslA of *Bacillus subtilis* (Bromley et al., 2015; Morris et al., 2017). Spider silk proteins also form β-sheet–rich elastic films at the air–water interface (Gustafsson et al., 2020; Tasiopoulos et al., 2021). Many recent studies have described the formation of elastic films by amyloid fibrils at oil–water interfaces in emulsions (reviewed in Xu et al., 2023). These examples suggest that the ability of proteins to self-assemble at interfaces and to form elastic films is deeply connected to their biological functions and offers opportunities for applications in biopolymer-based materials. A more detailed analysis of the structures adopted by RolA at interfaces will not only deepen our understanding of its biological roles but also shed light on the development of novel nanomaterials.

We also investigated the effects of both amorphous and rodlet films of RolA on the conidial surface properties of *A. oryzae* and found that surface hydrophobicity was significantly higher in conidia with rodlets than in those without (Figure 5G). This finding suggests that biological functions generally attributed to hydrophobins such as the hydrophobic modification of the conidial surface are expressed when RolA forms rodlets and that the amorphous state is functionally insufficient. The relationship between RolA structure and function identified *in vitro* is preserved on actual conidia.

Functional amyloids such as hydrophobins differ from pathogenic, misfolding-derived amyloids in that their structures are considered to be evolutionarily optimized to form fibrils efficiently (Sønderby et al., 2022). They self-assemble and form fibrils in a regulated manner and thereby contribute to morphogenesis rather than being toxic (Dragoš et al., 2017; Gebbink et al., 2005).

During fibril formation, amyloid proteins form an intermediate state, and these structures can sometimes be cytotoxic (Knowles et al., 2014). In the case of hydrophobins, amorphous films are formed at the air–water interface as intermediates (Takahashi et al., 2023; Terauchi et al., 2020) and might themselves have specific effects on fungal physiology. In this study, we demonstrated that the ability of RolA to express high surface activity and to fabricate mechanically robust films is achieved only in the rodlet state, whereas mutants that formed only an amorphous film lacked such properties (Figure 6). However, the immune evasion of *A. fumigatus* operates regardless of whether its hydrophobin is in the amorphous or rodlet state. Comparative proteomic analyses of the conidial surface covered by amorphous or rodlet films in *A. fumigatus* revealed substantial differences in the adsorbed proteins (Valsecchi et al., 2019). These findings suggest that interactions between hydrophobins and other proteins may be closely related to the structural state of hydrophobins. The regulation of fungal physiology mediated by hydrophobin self-assembly is essentially different from other biological regulatory mechanisms, making it especially important and intriguing.

**Figure 6.**
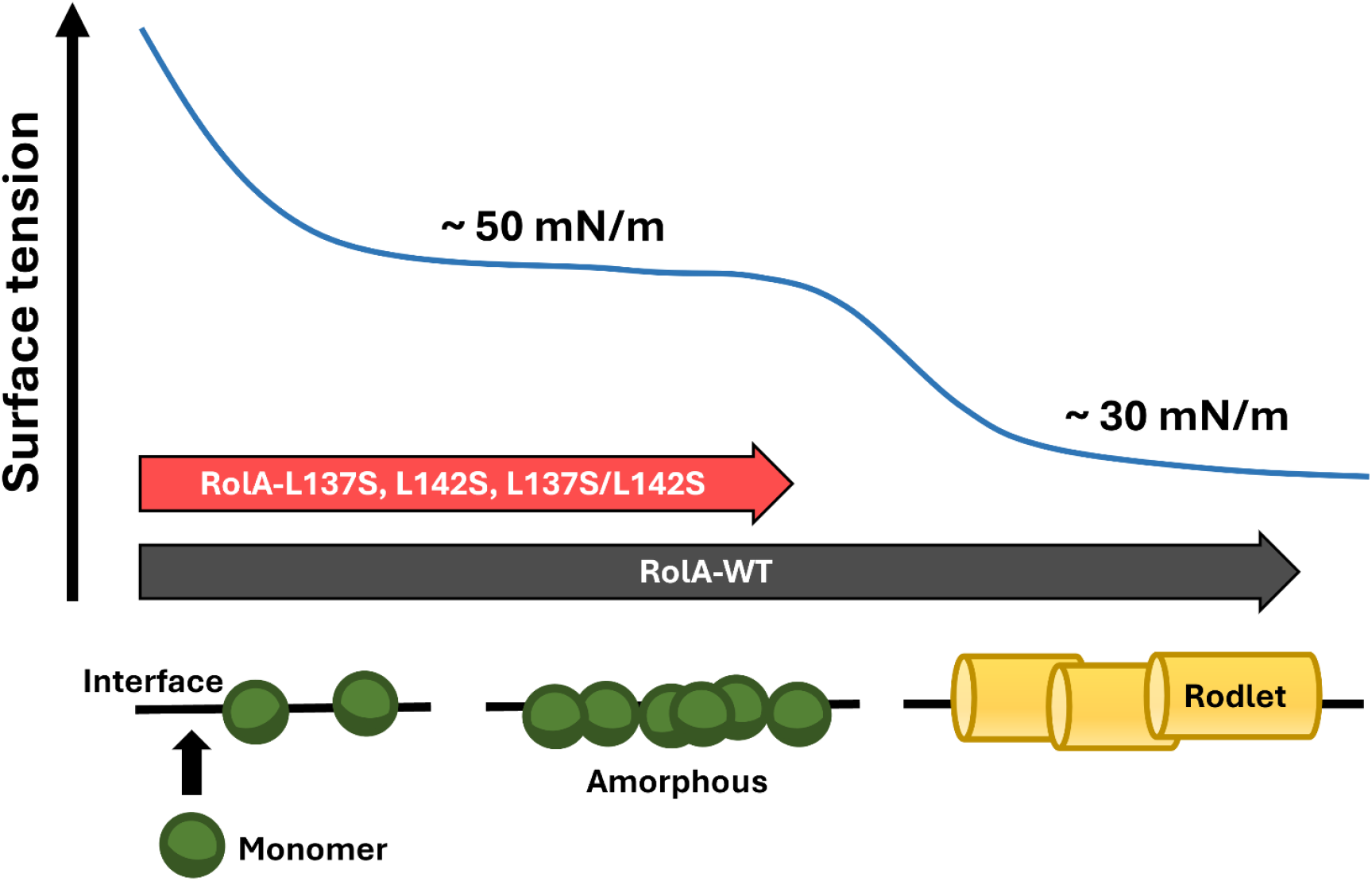
Schematic model of RolA adsorption and self-assembly. RolA first adsorbs to the interface and reaches an initial equilibrium interface tension, then self-assembles into rodlets to further reduce the interfacial tension and achieve a second equilibrium. In contrast to RolA-WT, RolA mutants that suppressed to form rodlets reach only the first equilibrium.

## 5 Conclusion

In this study, we used RolA-WT and mutants unable to form rodlets and clearly distinguished between amorphous and rodlet films of RolA to evaluate their surface properties in detail. We revealed that RolA forms a rigid film at the air–water interface and has high surface activity only when it is in the rodlet state. These properties directly influence the surface characteristics of *A.oryzae* conidia. These findings indicate that RolA self-assembly may substantially affect fungal physiology. The various other functionalities of hydrophobins should also be reconsidered from the perspective of structure–function relationships; understanding such relationships will be important for advancing hydrophobin research.

## 7 Author Contributions

D.I., N.T., and K.A. conceived and designed the research. D.I. and H.K. performed the experiments and D.I. and N.T. analyzed the data. D.I., N.T., Y.T., T.T., A.Y., K.M., M.M., H.Y., and K.A wrote the paper. K.A. supervised this project and funding acquisition. All authors discussed the results and commented on the manuscript. All authors have given approval to the final version of the manuscript.

## 9 Supplementary Material

The supplementary material for this article can be found online.

## 10 Data availability statement

The data underlying this article will be shared on reasonable request to the corresponding author.

