## Supporting Information for "Amorphous-to-Rodlet Structural Transition Governs the Interfacial Functions of *Aspergillus oryzae* Hydrophobin RolA"

Figure S1

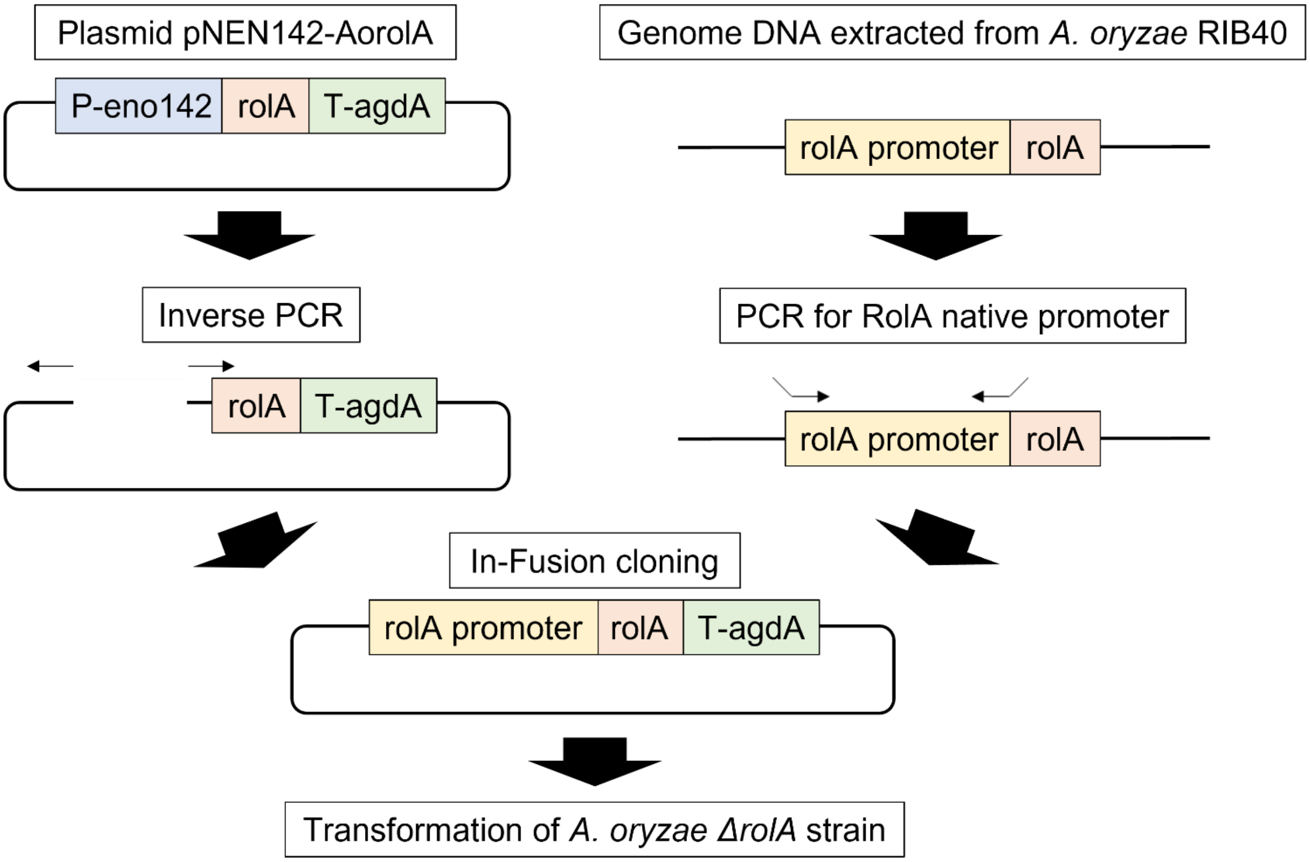


Construction of *A. oryzae* RolA-complemented strains. A plasmid expressing the RolA ORF under the control of the native promoter from the *A. oryzae* RIB40 strain was created and transformed into the *niaD* locus of the *ΔrolA* strain (parental strain: *adeA*⁻).

Figure S2

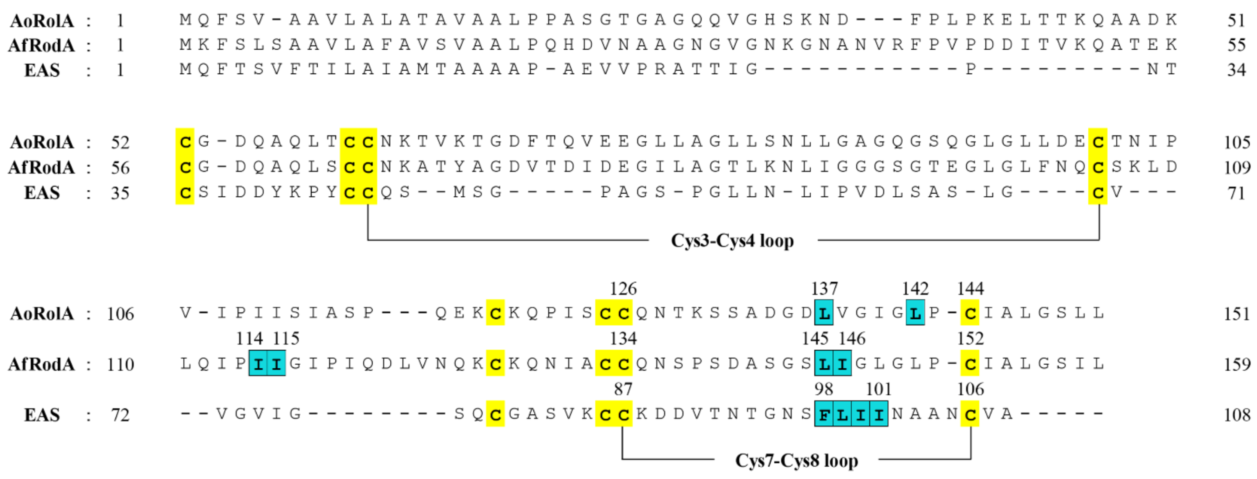


Alignment of the amino acid sequences of three Class I hydrophobins: *A. oryzae* RolA, *A. fumigatus* RodA, and *N. crassa* EAS. Cysteine residues that form conserved disulfide bonds are in yellow. Hydrophobic amino acid residues of RolA mutated in this study and those involved in rodlet formation in RodA and EAS are in cyan.

Figure S3

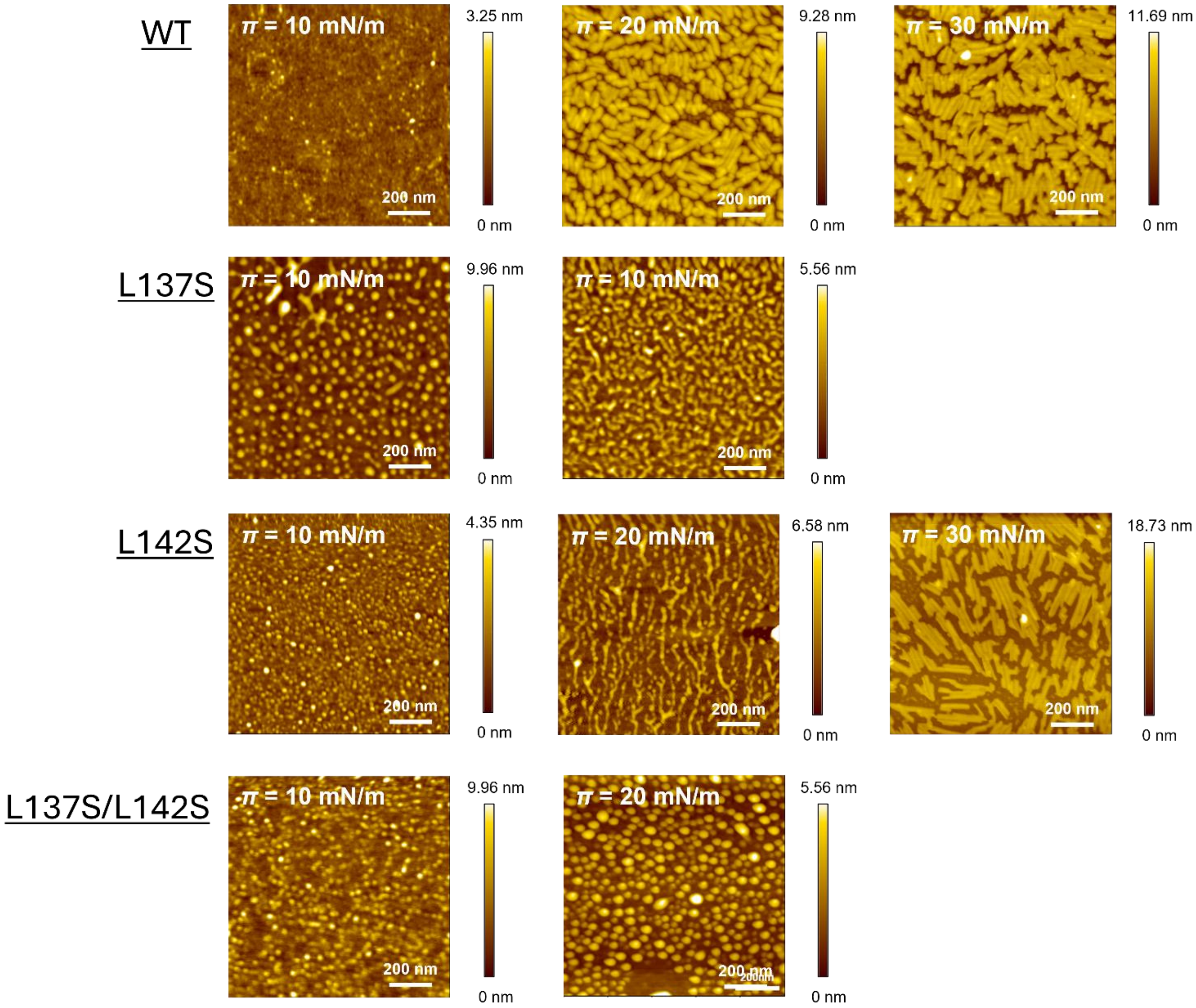


AFM topography images and height profiles of Langmuir films of wild-type RolA (WT) and its mutants. Films were transferred on hydrophobic silicon substrate at the indicated values of $\pi$. Image size, 1 μm × 1 μm.

Figure S4

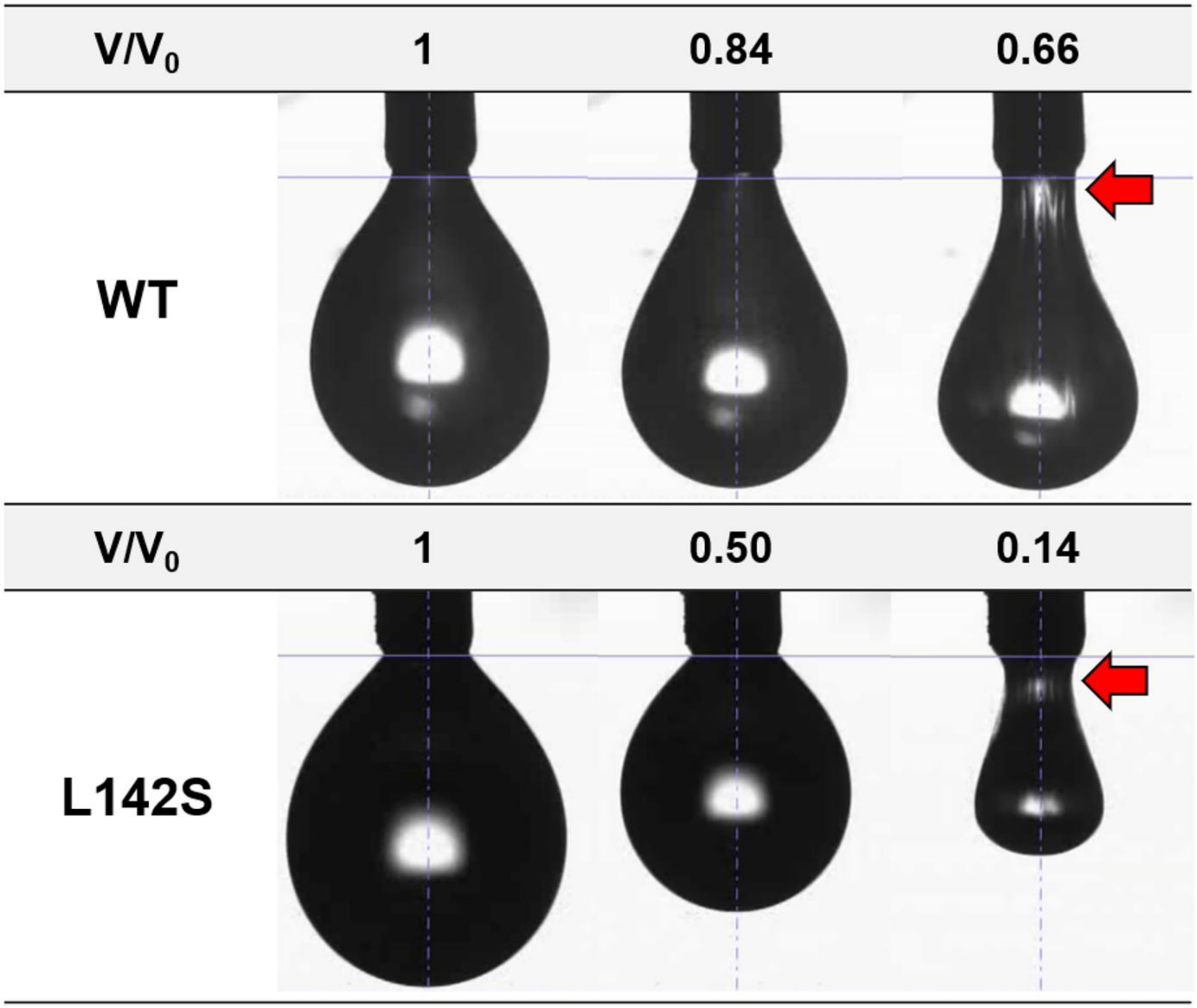


Compression ratios of RolA-WT and RolA-L142S droplets suspended in air. The concentration of RolA was 100 µg/ml. Clear buckling was observed in RolA-L142S droplets at compression ratios smaller than those of RolA-WT.

Figure S5

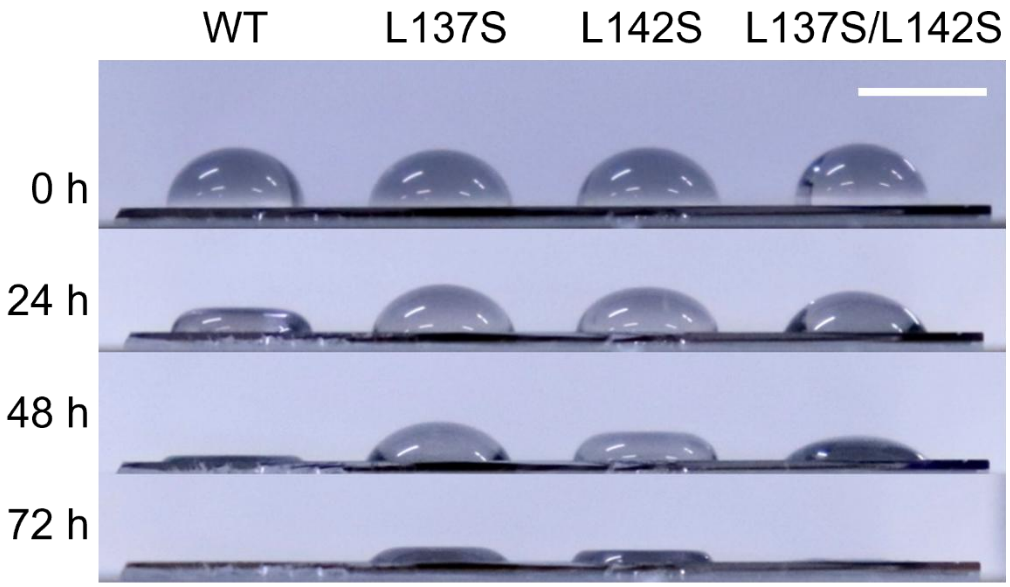


Flat droplets of the solutions of RolA-WT and its mutants. Plateaus were observed at the tops of all droplets. Scale bars = 2 mm.

Figure S6

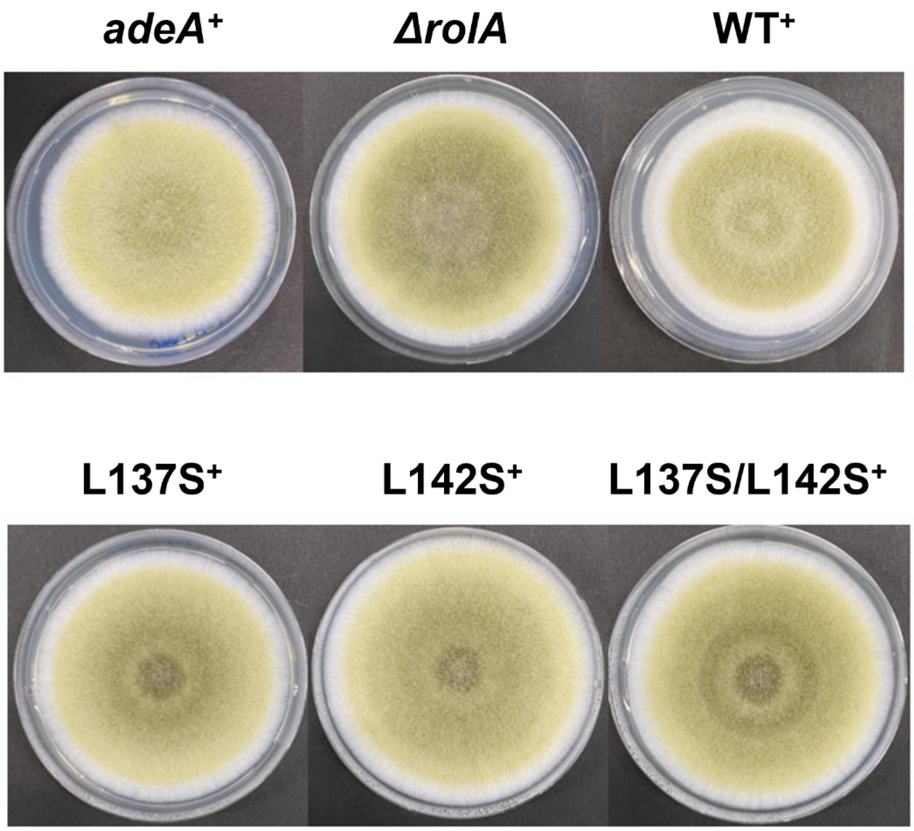


Colony morphology of *A. oryzae* control (*adeA*+), deletion (*ΔrolA*), and complemented strains on potato dextrose agar.

Table S1 Strains used in this study.

| Strain | Parent Strain | Genotype |
|---|---|---|
| RIB40 | | Wild-type strain (=ATCC42149) |
| eno-hyp WT | NSID-tApEnBdIVdV2 | *niaD*$^-$, *sC*$^-$, *adeA*$^-$, Δ*argB*::*adeA*$^-$, Δ*ligD*::*argB*, Δ*pyrG*::*adeA*, Δ*tppA*, Δ*pepE*, Δ*nptB*, Δ*dppIV*, Δ*dppV*::*pyrG*, *pNEN142-AorolA*::*niaD* |
| eno-hyp L137S | niaD300 (*niaD*$^-$) | *niaD*$^-$, *pNG-eno-hyp L137S*::*niaD* |
| eno-hyp L142S | niaD300 (*niaD*$^-$) | *niaD*$^-$, *pNG-eno-hyp L142S*::*niaD* |
| eno-hyp L137S/L142S | niaD300 (*niaD*$^-$) | *niaD*$^-$, *pNG-eno-hyp L137S/L142S*::*niaD* |
| *adeA*$^+$ | *adeA*$^-$ | *niaD*$^-$, *sC*$^-$, Δ*ligD*::*sC*, Δ*adeA*::*ptrA*, *adeA*$^+$ |
| Δ*rolA* | *adeA*$^-$ | *niaD*$^-$, *sC*$^-$, Δ*ligD*::*sC*, Δ*adeA*::*ptrA*, Δ*rolA*::*adeA* |
| RolA WT$^+$ | Δ*rolA* | *niaD*$^-$, *sC*-, Δ*ligD*::*sC*, Δ*adeA*::*ptrA*, Δ*rolA*::*adeA*, *rolA*$^+$, *RIB40p-AorolA WT*::*niaD* |
| RolA L137S$^+$ | Δ*rolA* | *niaD*$^-$, *sC*$^-$, Δ*ligD*::*sC*, Δ*adeA*::*ptrA*, Δ*rolA*::*adeA*, *rolA*$^+$, *RIB40p-AorolA L137S*::*niaD* |
| RolA L142S$^+$ | Δ*rolA* | *niaD*$^-$, *sC*$^-$, Δ*ligD*::*sC*, Δ*adeA*::*ptrA*, Δ*rolA*::*adeA*, *rolA*$^+$, *RIB40p-AorolA L142S*::*niaD* |
| RolA L137S/L142S$^+$ | Δ*rolA* | *niaD*$^-$, *sC*$^-$, Δ*ligD*::*sC*, Δ*adeA*::*ptrA*, Δ*rolA*::*adeA*, *rolA*$^+$, *RIB40p-AorolA L137S/L142S*::*niaD* |

Table S2 Primers used in this study.

| Primer name | Sequence (5’ to 3’) |
|---|---|
| L137S Fw | GGCGACTCCGTCGGTATTGGTCTTCC |
| L137S Rv | CCGACGGAGTCGCCATCCTGTGATTG |
| L142S Fw | ATTGGTTCCCCTTGCATCGCTCTCGGC |
| L142S Rv | GCAAGGGGAACCAATACCGACGAGGTC |
| L137S/L142S Fw | GGCGACTCCGTCGGTATTGGTTCCCC |
| L137S/L142S Rv | ACCGACGGAGTCGCCATCCTGTGATTG |
| pNEN142-enoA-RolA Fw1 | TCTTCGCTATTACGCCAGCTG |
| pNEN142-enoA-RolA Fw2 | CCCATAGGTGAGTTTGGTTG |
| pNEN142-enoA-RolA Fw3 | GTAGGGATATCATCGCATGGC |
| pNEN142-enoA-RolA Fw4 | CCTCCCTTTCCGTCTCTTTTC |
| pNEN142-enoA-RolA Rv1 | GGACGAGGTCCAACACTCAC |
| pNEN142-enoA-RolA Rv2 | ATTTGGTCGATGAAGACGACTC |
| pNEN142-enoA-RolA Rv3 | GCAACTCGCTTACCGATTACG |
| pNEN142-enoA-RolA Rv4 | CCCCGTAGAATCACGAATGAG |
| In-Fusion Fw2 | ATCTTCCGGTATGGCTGTTG |
| In-Fusion Fw3 | GAAGCAGTGAGATGGATGAGG |
| In-Fusion Fw4 | CCAGGGTCTTGGTCTCTTGG |
| In-Fusion Rv1 | GGACGAGGTCCAACACTCAC |
| In-Fusion Rv2 | ATTTGGTCGATGAAGACGACTC |
| In-Fusion genome Fw | AAACGACGGCCAGTGGGACCGTGCAGTAGTAGAGTTC |
| In-Fusion genome Rv | ATGCCATATGACTAGGTGATGGTGGCTTTGGTTTTG |
| In-Fusion InversePCR Fw | CTAGTCATATGGCATGCAGTTC |
| In-Fusion InversePCR Rv | CACTGGCCGTCGTTTTACAAC |
| AorolA-top-f | GGACCGTGCAGTAGTAGAGTTCCTAAC |
| AorolA-top-r | CGCGTGCGCAGGAGTGTTTGTTGTGTGATGGTGGCTTTGGTTTTGAACGAG |
| AnAdeA-f | CAACAAACACTCCTGCGCACGCG |
| AnAdeA-r | GGTACCTGATGGCCTCGCAGATAAA |
| AorolA-bottom-f | GTTTATCTGCGAGGCCATCAGGTACCGCGATTGCATTCGCGAAAATGGTAGCTC |
| AorolA-bottom-r | GTTCGGGGTTTTTTCTTTGGTGTGTACTTGGTGC |
| AoRolA-delta-L-chk-f2 | TACCAAACGCGGGCACCAG |
| AoRolA-delta-R-chk-r2 | AATCGGCGGATGAGTCG |
| AnAdeA-chk-f2 | GGTAAACGAGCCGAGAGAATC |
| AnAdeA-chk-r2 | GGTAGCAGAATTGTCTCCCATC |
| AoRolA-chk-f | ACCACCAAGCAGGCCG |
| AoRolA-chk-r | GAGCGATGCAAGGAAGAC |
| Ao Histone-RT Fw | ACGTCGTCTATGCCCTCAA |
| Ao Histone-RT Rv | GGAAGGACAACCGACGCG |
| Ao kexB-RT Fw | GCGACATCAGTGTGGAGTTG |
| Ao kexB-RT Rv | GACCGTCCACTTTCCAACAC |
| Ao RolA-RT Fw | TGCACCAACATCCCTGTTATCC |
| Ao RolA-RT Rv | GGACTTGGTGTTCTGGCAGC |

Table S3 Height, width, and length of rodlets and rod-like structures formed by hydrophobin RolA. All numerical values are in nanometers.

| **Substrate** | **RolA** | **$\pi$ = 20 mN/m** | | | **$\pi$ = 30 mN/m** | | | **$\pi$ = 40 mN/m** | | |
|---|---|---|---|---|---|---|---|---|---|---|
| | | **Height** | **Width** | **Length** | **Height** | **Width** | **Length** | **Height** | **Width** | **Length** |
| Hydrophilic silicon | WT | 2.63 ± 0.15 | 12.0 ± 1.83 | 76.0 ± 34.2 | 2.10 ± 0.40 | 12.0 ± 1.83 | 95.9 ± 35.0 | – | | |
| | L137S | 2.70 ± 0.33 | 12.4 ± 3.05 | 73.7 ± 38.5 | – | | | – | | |
| | L142S | No rodlets | | | 2.78 ± 0.32 | 11.6 ± 2.10 | 93.0 ± 41.1 | 3.22 ± 0.21 | 11.5 ± 1.91 | 76.2 ± 31.1 |
| | L137S/L142S | No rodlets | | | – | | | – | | |
| Hydrophobic silicon | WT | 6.85 ± 0.36 | 19.9 ± 3.58 | 69.6 ± 26.3 | 7.96 ± 0.40 | 20.5 ± 2.11 | 89.4 ± 25.7 | – | | |
| | L137S | No rodlets | | | – | | | – | | |
| | L142S | No rodlets | | | 6.87 ± 0.83 | 16.0 ± 2.54 | 96.4 ± 48.0 | 8.27 ± 0.67 | 20.5 ± 2.74 | 57.9 ± 19.6 |
| | L137S/L142S | No rodlets | | | – | | | – | | |

Table S4

Water contact angle of RolA Langmuir film transferred on the hydrophobic substrate

| **RolA** | **Without RolA** | **Surface pressure** | | |
|---|---|---|---|---|
| | | ***π* = 10 mN/m** | ***π* = 20 mN/m** | ***π* = 30 mN/m** |
| WT | 99.6 ± 1.0° | 76.8° ± 0.4° | 52.2° ± 2.6° | 39.1° ± 3.7° |
| L137S | | 85.1° ± 2.0° | 75.7° ± 0.8° | – |
| L142S | | 83.1° ± 0.4° | 80.4° ± 0.3° | 78.3° ± 0.6° |
| L137S/L142S | | 88.9° ± 0.3° | 75.9° ± 1.2° | – |

All data are shown as means ± standard deviation ($n \geq 3$).
